# Interrogating Metabolic Interactions Between Skeletal Muscle and Liver Circadian Clocks In Vivo

**DOI:** 10.1101/2022.02.27.482160

**Authors:** Jacob G. Smith, Kevin B. Koronowski, Tomoki Sato, Carolina Greco, Paul Petrus, Amandine Verlande, Siwei Chen, Muntaha Samad, Ekaterina Deyneka, Lavina Mathur, Ronnie Blazev, Jeffrey Molendijk, Thomas Mortimer, Arun Kumar, Oleg Deryagin, Mireia Vaca-Dempere, Peng Liu, Valerio Orlando, Benjamin L. Parker, Pierre Baldi, Patrick-Simon Welz, Cholsoon Jang, Selma Masri, Salvador Aznar Benitah, Pura Muñoz-Cánoves, Paolo Sassone-Corsi

**Affiliations:** Center for Epigenetics and Metabolism, U1233 INSERM, Department of Biological Chemistry, University of California, Irvine, CA, 92697, USA; Department of Experimental and Health Sciences, Pompeu Fabra University (UPF), Parc de Recerca Biomèdica de Barcelona (PRBB), Barcelona, 08003, Spain; Department of Biochemistry and Structural Biology, Joe R. & Teresa Lozano Long School of Medicine, Sam and Ann Barshop Institute for Longevity and Aging Studies, University of Texas Health San Antonio, San Antonio, Texas, 78229, USA; Laboratory of Nutritional Biochemistry, Graduate School of Nutritional and Environmental Sciences, University of Shizuoka, Shizuoka 422-8526, Japan; Department of Biomedical Sciences, Humanitas University and Humanitas Research Hospital IRCCS, Via Manzoni 56, 20089, Rozzano (Milan), Italy; Department of Biological Chemistry, Center for Epigenetics and Metabolism, Chao Family Comprehensive Cancer Center, University of California, Irvine, Irvine, CA, 92697, USA; Institute for Genomics and Bioinformatics, Department of Computer Science, University of California, Irvine, CA, 92697, USA; Department of Anatomy and Physiology, The University of Melbourne, VIC 3010, Australia; Institute for Research in Biomedicine (IRB Barcelona), The Barcelona Institute of Science and Technology (BIST), Barcelona, 08028, Spain; King Abdullah University of Science and Technology, KAUST Environmental Epigenetics Research Program, Biological and Environmental Sciences and Engineering Division, Thuwal 23955, Saudi Arabia; Program in Cancer Research, Hospital del Mar Medical Research Institute (IMIM), Parc de Recerca Biomèdica de Barcelona (PRBB), Barcelona, 08003, Spain; Catalan Institution for Research and Advanced Studies (ICREA), Barcelona, 08010, Spain; Spanish National Center for Cardiovascular Research (CNIC), Madrid, 28029, Spain

## Abstract

Expressed throughout the body, the circadian clock system achieves daily metabolic homeostasis at every level of physiology, with clock disruption associated with metabolic disease (*1*, *2*). Molecular clocks present in the brain, liver, adipose, pancreas and skeletal muscle each contribute to glucose homeostasis (*3*). However, it is unclear; 1) which organ clocks provide the most essential contributions, and 2) if these contributions depend on inter-organ communication. We recently showed that the liver clock alone is insufficient for most aspects of daily liver glucose handling and requires connections with other clocks (*4*). Considering the pathways that link glucose metabolism between liver and skeletal muscle, we sought to test whether a clock connection along this axis is important. Using our previous published methodology for tissue-specific rescue of *Bmal1* in vivo (*4*, *5*), we now show that in the absence of feeding-fasting cycles, liver and muscle clocks are not sufficient for systemic glucose metabolism, nor do they form a functional connection influencing local glucose handling or daily transcriptional rhythms in each tissue. However, the introduction of a daily feeding-fasting rhythm enables a synergistic state between liver and muscle clocks that leads to restoration of systemic glucose tolerance. These findings reveal limited autonomous capabilities of liver and muscle clocks and highlight the need for inter-organ clock communication for glucose homeostasis which involves at least two peripheral metabolic organs.

## Independence of liver and muscle autonomous clocks in vivo

To determine both the local roles of and interactions between autonomous liver and muscle clocks, we used *Bmal1-*stopFL mice, wherein endogenous expression of the critical clock component *Bmal1* (*6*) can be reconstituted exclusively in Cre recombinase-expressing tissues in otherwise *Bmal1*-deficient animals (*5*). *Bmal1-* stopFL mice were crossed with hepatocyte-driven Alfp-Cre and skeletal muscle-driven Hsa-Cre lines to generate the following genotypes: wild-type (WT, all clocks); full *Bmal1* knockout (KO, no clocks); liver *Bmal1*-reconstituted (Liver-RE, only liver clock) (*4*, *5*); skeletal muscle *Bmal1*-reconstituted (Muscle-RE, only skeletal muscle clock); both liver and skeletal muscle *Bmal1*-reconstituted (Liver+Muscle-RE, only liver and skeletal muscle clocks) (Fig. 1a). Western blotting confirmed expected tissue-specificity of BMAL1 expression across genotypes (Extended Data Fig. 1a–d).

**Fig. 1:**
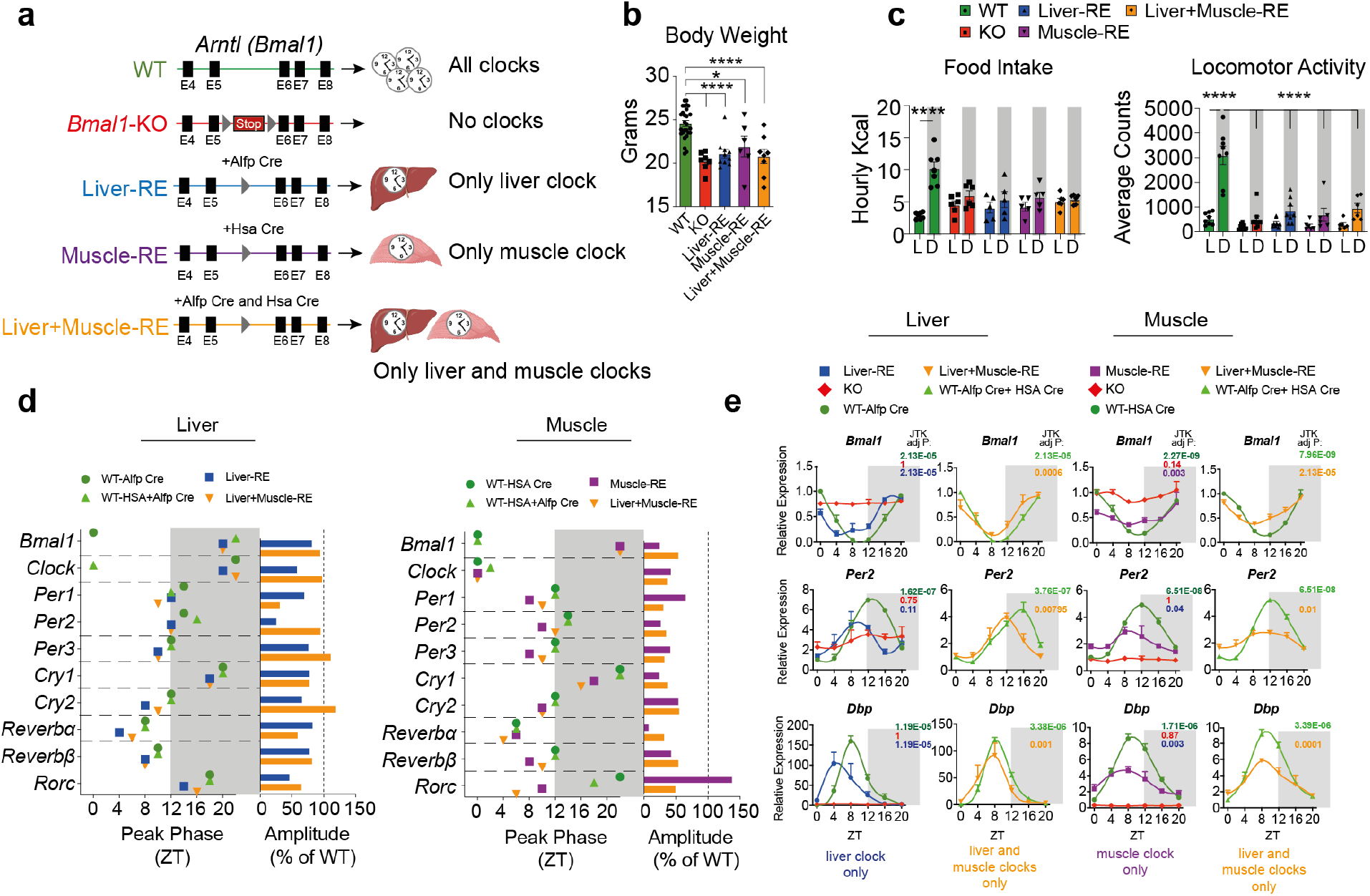
Liver and muscle clocks oscillate in a tissue-autonomous manner. **a**, genetic scheme for tissue-specific clock reconstitution (RE) in mice. **b**, body weight at 8 weeks old left (n= 6-28) * p<0.05, ****p<0.0001 by one way analysis of variance (ANOVA), with Fisher’s LSD test, **c**, food intake of RE mice under 12h light:12h dark (L:D) conditions, two-way ANOVA with Bonferroni post-test, **** p<0.0001, n=5-7. Right, locomotor activity, two-way ANOVA with Bonferroni post-test, **** p<0.0001, n=6-9. **d-e**, RNA sequencing of livers and gastrocnemius muscles harvested around the clock under 12 hr light: 12 hr dark conditions (n=3). Liver sequencing data for WT-Alfp Cre, KO and Liver-RE from (*8*). No full-length *Bmal1* mRNA or protein of any size is detected in KO mice-see Extended Data Figure 1a–d. **d**, peak phase and amplitude of clock genes from JTK_CYCLE rhythmicity analysis. **e**, clock gene expression in liver and muscle of RE mice, JTK_CYCLE Adjusted P values noted for each genotype.

Although total food intake was similar across all genotypes (Extended Data Fig. 1e), KO mice exhibited lower body weight and higher fat to lean mass ratio versus age-matched WT animals, and the presence of liver and/or muscle clocks did not alter either parameter further (Fig. 1b, Extended Data Fig. 1f). Due to loss of brain clock function (*7*), and as reported previously for Liver-RE mice (*4*, *8*, *9*), KO and RE mice lacked robust 24h-rhythms of food intake and displayed severely blunted dark phase locomotor activity under 12h light: 12 dark conditions and *ad libitum* feeding (Fig. 1c). As a result, only WT mice had consistent and robust diurnal metabolic cycles as assayed through respiratory exchange ratio and energy expenditure (Extended Data Fig. 1g,h). Just 6 of 23 KO and RE mice tested had significant rhythms of both parameters (Extended Data Fig. 1g), and the presence of both liver and muscle clocks had no additive effect (Extended Data Fig. 1g,h).

Using Liver-RE mice, we previously demonstrated the liver core clock is partially autonomous, oscillating with reduced amplitude and a ~2-4 hour phase advance versus WT (*4*, *8*). To assess muscle clock autonomy, and probe potential muscle-liver connections, we performed RNA-Sequencing on gastrocnemius muscles (predominantly fast-twitch fibers (*10*)) and livers harvested every 4-hours over the diurnal cycle. As expected, loss of *Bmal1* resulted in loss of clock gene oscillations (Extended Data Fig. 2). In Muscle-RE, similar to Liver-RE, the clock genes *Bmal1*, *Per2* and *Dbp* oscillated with reduced amplitude and were phase advanced versus WT (Fig. 1d–e). Overall clock gene oscillations were considerably more pronounced in liver, revealing a less robust autonomous clock in muscle and highlighting tissue-specific differences in autonomous clock function (Fig. 1d–e, Extended Data Fig. 2). Oscillations were remarkably similar in muscles and livers of all RE genotypes, though peak phases in liver were modestly corrected for certain genes, including *Dbp* and *Reverbα* (Fig. 1d,e, Extended Data Fig. 2). These findings demonstrate that the autonomous muscle clock exhibits reduced amplitude and robustness versus the liver clock, and that liver and muscle autonomous clocks oscillate independently of one another in vivo.

Next, we analyzed full diurnal transcriptomes to assess whether genome-wide rhythmic transcription shows evidence of muscle clock autonomous output, as well as synergy between liver and muscle autonomous clocks. For Liver-RE and its controls, we used data from our previous study which featured mice of identical age, sex, diet and conditions (*8*). As expected, loss of *Bmal1* resulted in loss of most oscillating transcripts (Extended Data Fig. 3a). By JTK_CYCLE, the number of oscillating transcripts in the livers of Liver-RE and Liver+Muscle-RE was similar; 654 in Liver+Muscle-RE, 666 in Liver-RE. Compared to respective WTs, 12.39% of rhythmic hepatic transcription was restored in Liver+Muscle-RE versus 12.11% in Liver-RE (Fig. 2a and Extended Data Fig. 3a). This was accompanied by improvements in peak phase compared to WT (Fig. 2b, Extended Data Fig. 3b), however amplitudes remained dampened compared to WT (Extended Data Fig. 3b, Fig. 2c). As expected, liver oscillating transcripts were enriched similarly for circadian rhythm and metabolic pathways in single and double RE livers (Fig. 2d). Representative genes from these pathways showed similar expression patterns in livers of Liver-RE and Liver+Muscle-RE (Fig. 2e). We also found minimal differences between Liver-RE and Liver+Muscle-RE oscillating transcriptomes upon interrogation of our data via BIO_CYCLE analysis, which uses an alternative algorithmic approach to define circadian-regulated transcripts (Extended Data Fig. 4). Probing muscle output autonomy and influence in the other direction (from liver to muscle), we found Muscle-RE mice were only able to recapitulate 4.05% of WT oscillating transcripts, well below the liver’s ability of 12.11% (Fig. 2a). This is consistent with a less robust autonomous core clock in muscle (Fig. 1d,e, Extended Data Fig. 1a–d, Extended Data Fig. 2). A comparable number of oscillating transcripts were detected in muscles of Muscle-RE and Liver+Muscle-RE (155 and 170) and overlap with respective WTs (4.05% and 3.95%) was also comparable (Fig. 2a and Extended Data Fig. 3a,b). Like liver, amplitudes in muscle remained dampened yet changes in peak phase were observable for some oscillating transcripts with both clocks present (Extended Data Fig. 3a,b, Fig 2b–c). In single and double RE muscles, oscillating transcripts were enriched for circadian rhythm, and enrichments were otherwise limited due to the small number of oscillating transcripts (Fig. 2d). Assessment of key metabolic genes in muscle revealed similar oscillatory profiles in Muscle-RE and Liver+Muscle-RE mice (Fig. 2e). BIO_CYCLE analysis also failed to uncover a synergy between liver and muscle clocks (Extended Data Fig. 4) and autonomous output was highly tissue-specific (Extended Data Fig. 3c, 4c). Our results highlight the independent nature of liver and muscle autonomous clock function and transcriptional output in vivo, highlighting the importance of inputs from the broader clock network for the full repertoire of circadian rhythms in each tissue.

**Fig. 2:**
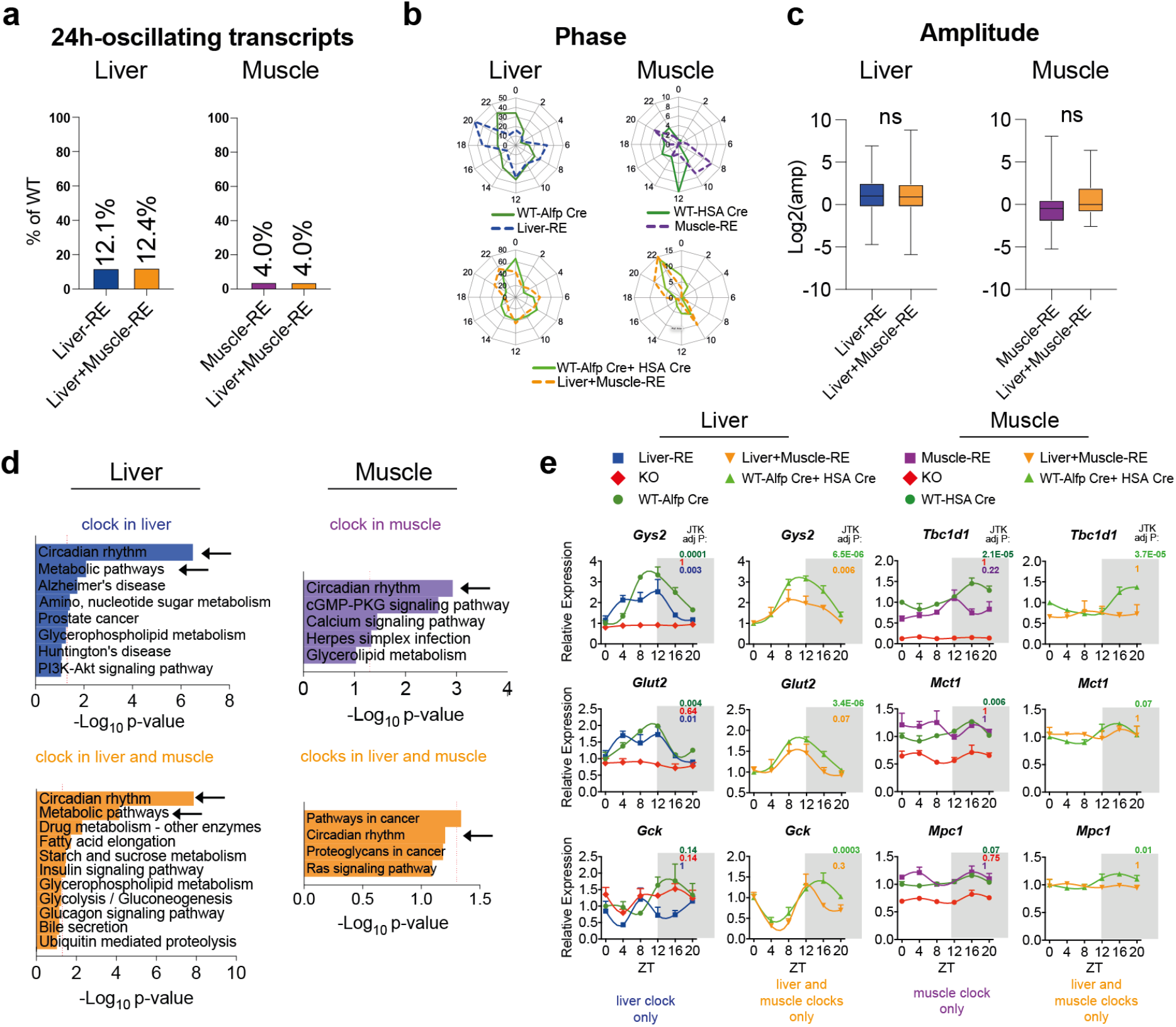
Independence of liver and muscle autonomous clock transcriptional output. **a-e**, RNA sequencing of livers and gastrocnemius muscles harvested around the clock under 12 hr light: 12 hr dark conditions (n=3) with subsequent JTK_CYCLE rhythmicity analysis for transcripts oscillating with a 24h period. Liver sequencing data for WT-Alfp Cre and Liver-RE from (*8*). **a**, number of oscillating transcripts expressed as percentage restoration of wild type. **b**, peak phase, and amplitude. **c**, of 24h rhythmic transcripts in RE mice, only transcripts that also oscillate in WT are included. For amplitude comparisons within tissue, ns= not significant by two-tailed t-test, data displayed as box and whiskers min to max, line at median **d**, gene set enrichment analysis with KEGG of 24h-rhythmic transcripts shared between WT and denoted RE mice, arrows indicate pathways present in single and double RE mice within each tissue. **e**, expression of metabolism-related transcripts.

## Insufficiency of liver and muscle autonomous clocks for aspects of glucose metabolism

Liver and skeletal muscle are the major organs that achieve whole-body glucose homeostasis (*3*). Muscle is the primary site of postprandial glucose disposal and liver is known to take up muscle-derived metabolites of glucose (e.g. lactate liberated during the Cori cycle (*11*)). Knowing from previous studies that BMAL1 is necessary for muscle glucose uptake (*12*, *13*), we asked whether it is also sufficient. To follow the metabolism of glucose into tissues and assess its conversion into other metabolites, we performed isotope tracing with ^13^C-labeled glucose. Following a 6 hour fast, mice were gavaged with uniformly labeled ^13^C glucose at 1.5 g/kg dose (typical dose for glucose tolerance test), which achieved sufficient serum glucose labeling (~ 50%) across all genotypes for downstream metabolite measurements (Extended Data Fig. 5a). Tissues were collected 25-30 min post gavage to capture the peak labeling of downstream metabolites in liver and skeletal muscle (*14*) by liquid chromatography-mass spectrometry (Fig. 3a).

**Fig. 3:**
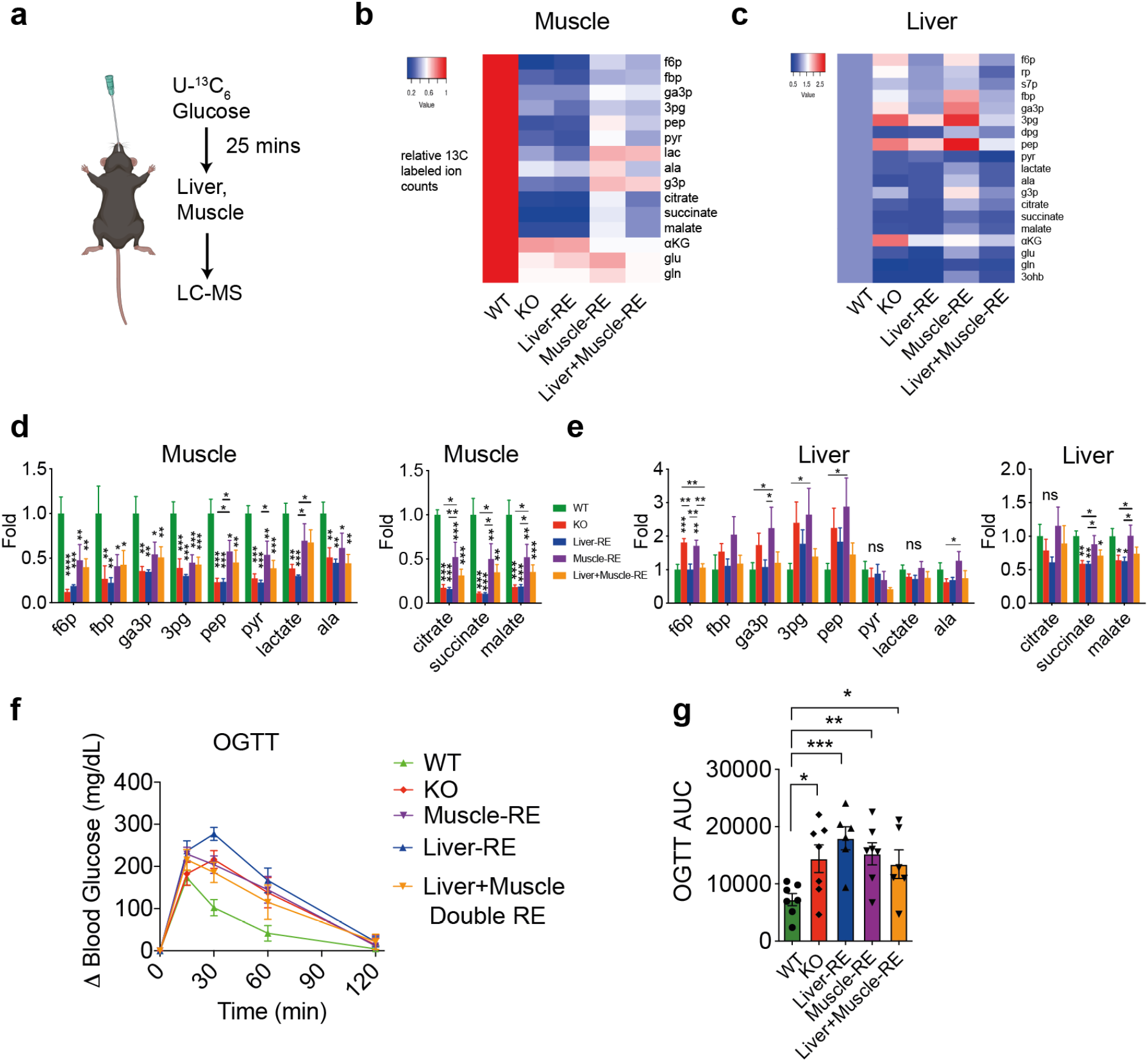
Partial control of local but not systemic glucose metabolism by liver and muscle autonomous clocks under ad libitum conditions. **a**, isotope tracing experimental design. Mice were fasted for 6 hr from ZT0 to ZT6, during a WT mouse’s normal fasting period. 1.5 g/kg of uniformly labeled glucose (U-^13^C_6_) was gavaged orally and tissues were harvested 25-30 min later. Metabolite labeling was measured by LC-MS and normalized to circulating glucose labeling (n=5-6). Data are presented as ^13^C-labeled ion counts relative to WT. Heatmaps from muscle **b**, and liver **c**, showing incorporation of glucose-derived ^13^C labelled carbons in the indicated metabolites. f6p, fructose 6-phosphate; fbp, fructose 1,6-bisphosphate; rp, ribose phosphate; s7p, sedoheptulose 7-phosphate; ga3p, glyceraldehyde 3-phosphate; 3pg, 3-phosphoglycerate; dpg, 2,3-diphosphoglycerate; pep, phosphoenolpyruvate; pyr, pyruvate; lac, lactate; ala, alanine; g3p, glycerol 3-phosphate; KG, alpha-ketoglutarate; glu, glutamate; gln, glutamine. 3ohb, 3-hydroxybutyrate. **d-e**, left – labeling of glycolysis metabolites; right – labeling of TCA cycle metabolites; * p<0.05, ** p<0.01, *** p<0.001, ****p<0.0001 by one way analysis of variance (ANOVA), with Fisher’s LSD test, performed for each metabolite. **f**, oral glucose tolerance tests (OGTT) with unlabeled glucose were performed as described in (**a**) (n=6-7) and are expressed as delta change from baseline blood glucose. **g**, area under the curve (AUC) for OGTTs, ns= not significant, * p<0.05, ** p<0.01, *** p<0.001, one-way ANOVA with Fishers LSD test.

Compared to WT muscles, KO muscles showed substantially lower labeling of glycolytic and TCA cycle intermediates, indicating impaired glucose uptake and oxidation (Fig 3b,d). The presence of muscle BMAL1 in Muscle-RE mice markedly restored labeling of these metabolites, but values were still considerably lower than WT, highlighting the requirement of other tissue clocks for muscle glucose metabolism. The restoration of the liver clock was not sufficient to improve muscle glucose metabolism, as Muscle-RE and Liver+Muscle-RE labeling patterns were similar (Fig. 3b,d). Liver is also known to take up glucose during feeding, to support anabolic pathways (e.g., glycogen and fat synthesis), as well as take up muscle-derived metabolites of glucose (e.g., lactate and alanine) for oxidation or recycling (*15*). In contrast to muscles, KO livers displayed increased labeling of glycolytic intermediates yet reduced labeling of TCA cycle intermediates (Fig. 3c,e). In Liver-RE mice, we observed normalized labeling of glycolytic but not TCA cycle intermediates, suggesting disparate regulation mechanisms. Intriguingly, Muscle-RE livers exhibited increased labeling of TCA cycle intermediates compared to KO, suggesting muscle-liver metabolic crosstalk in the form such of liver processing of muscle-clock dependent, glucose-derived metabolites. This phenotype was not evident in Liver+Muscle-RE mice, likely indicating that the liver clock dominates over the muscle clock for regulating its own TCA cycle (Fig. 3c,e). Taken together, these data show partial rescue of tissue glucose metabolism by autonomous clocks in muscle and liver, with no evidence of further improved metabolism with both clocks present.

In addition to maintaining metabolic homeostasis within the tissue, liver and muscle glucose metabolism helps balance the consumption and production of glucose over the course of a day to maintain blood glucose levels within a physiological range (*16*, *17*). To ask whether any RE mice have better glycemic control than KO, we performed oral glucose tolerance tests. Compared to WT, glucose tolerance was impaired in all other genotypes (Fig. 3f,g) in both genders (Extended Data Fig. 5b,c). Even though intratissue glucose metabolism is partially rescued by local BMAL1, our OGTT results demonstrate the insufficiency of BMAL1 in either liver or muscle for systemic glucose homeostasis, as well as the lack of functional compensation between liver and muscle clocks in this context.

## Feeding rhythms bolster autonomous liver and muscle clocks, enabling synergistic control of glucose tolerance

The observed glucose intolerance of Liver+Muscle-RE mice under ad libitum conditions prompted the question of which key circadian signals are absent in these animals. KO and RE mice lack a feeding-fasting rhythm, a brain clock-driven circadian behavior that both synchronizes and interacts with local clocks to control transcript oscillations in peripheral organs (*7*, *8*, *18*–*22*). We previously demonstrated that feeding-fasting behavior is integrated by the autonomous liver clock, an interplay that bolsters the phase and output of the clock as well as improves daily glucose pathways within liver (*8*). If feeding behavior is integrated similarly in muscle, it is plausible that a connection might emerge between bolstered liver and muscle clocks, allowing proper regulation of systemic glucose metabolism. To this end, we performed experiments under night feeding conditions, wherein mice only had access to food during the 12 hour dark period (ZT12-ZT24/0) for 2 weeks (Fig. 4a). After 2 weeks of night feeding, only small changes in body weight were observed (WT = +2.14%; KO and RE = −3.48% to −6.81%) (Extended Data Fig 6a). Night feeding induced a feeding-fasting rhythm in KO and RE mice, as well as an accompanying rhythm of serum insulin, evidenced by a peak during feeding at ZT16 and trough during fasting at ZT4 (Fig. 4b,c). We have previously demonstrated serum insulin rhythms under night feeding in KO and Liver-RE mice (*8*). A rhythm of serum insulin is noteworthy because it can act as a systemic synchronizing cue for circadian clock function in peripheral organs (*23*, *24*).

**Fig. 4:**
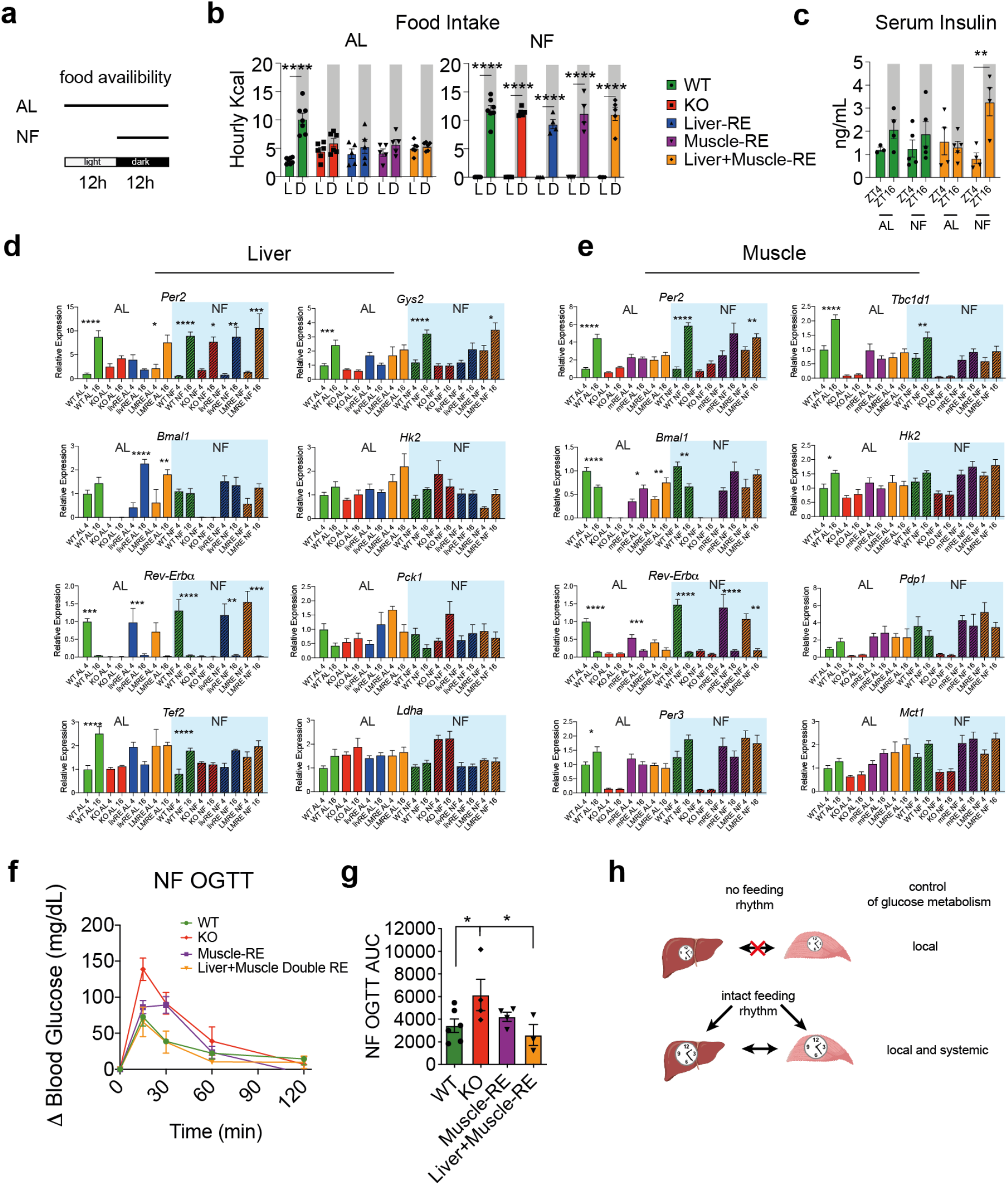
Muscle-liver clock synergy under night feeding mediates systemic glucose tolerance. **a**, experimental setup for night fed (NF) versus ad libitum (AL) fed mice. **b**, food intake in AL vs NF mice, **** p< 0.0001 by two-way analysis of variance (ANOVA), with Bonferroni post-test, n=4-7; AL panel from Figure 1, included for clarity. **c**, serum insulin in WT and Liver+Muscle-RE mice under AL and NF conditions, ** p< 0.01 by two-way analysis of variance (ANOVA), with Bonferroni post-test, n=4-5. **d-e**, gene expression from qPCR in liver and muscle of clock and metabolic genes at ZT4 and 16 under AL and NF conditions, n=3-6, * p<0.05, ** p<0.01, *** p<0.001, ****p<0.0001 by two-way ANOVA with Bonferroni post-test. **f**, oral glucose tolerance test (OGTT) in NF conditions expressed as delta change from baseline blood glucose, n=2-6. **g**, corresponding area under the curve (AUC) for NF OGTT, * p<0.05, ** p<0.01, *** p<0.001, by one-way ANOVA, with Fishers LSD test. **h**, schematic of the minimal clock network for required for local and systemic glucose tolerance.

Next, we analyzed the transcriptional response of each genotype to the *de novo* feeding-fasting rhythm. We measured expression of clock and metabolic genes at ZT4 and ZT16, coinciding with serum measurements. Under night feeding, whereas livers without *Bmal1* (KO and Muscle-RE) displayed markedly lower or mistimed gene expression, livers expressing BMAL1 (Liver-RE and Liver+Muscle-RE) displayed the most WT-like gene expression under night feeding (Fig. 4d, Extended Data Fig. 6b). This was true for clock genes (e.g. *Bmal1*, *Reverbα*, *Tef*, *Hlf*) and glucose metabolism genes (e.g. *Gys2*, *Hk2*, *Ldha*, *Pck1*) (Fig. 4d). Thus, as we showed previously, the liver clock is required to transcriptionally integrate the feeding-fasting rhythm (*8*). Certain liver genes (*Reverbα*, *Gys2*) appeared to benefit from muscle BMAL1 expression in Liver+Muscle-RE mice, but effects were modest (Fig. 4d, Extended Data Fig 6b).

In muscle, both *Per2* and *Reverbα* oscillated weakly under *ad libitum* conditions, however night feeding normalized expression to WT in muscles expressing BMAL1 (Muscle-RE and Liver+Muscle-RE) but not KO (Fig. 4e). Thus, *Per2* and *Reverbα* oscillations are primarily driven by the integration of the autonomous muscle clock and feeding-fasting rhythm. Other clock genes such as *Per3* displayed corrections of their levels of expression under night feeding (Extended Data Fig. 6c). Expression of key glucose metabolism genes, including *Tbc1d1*, *Hk2*, *Pdp1* required both the autonomous muscle clock and feeding-fasting rhythm (Fig. 4e), revealing integration of feeding behavior by the local clock also occurs in muscle. Finally, for all genes tested, the effects of liver BMAL1 on muscle gene expression were modest if at all present (Fig. 4e, Extended Data Fig. 6c). Together, these data demonstrate autonomous clock integration of daily feeding-fasting behavior bolsters clock and metabolic gene expression in both liver and muscle. Because single and double RE responses to night feeding were similar, this integration is occurring independently in both liver and muscle and is not overtly influenced by the presence of the other clock.

We next asked whether feeding-rhythm-bolstered autonomous clock function would improve serum glucose responses, specifically in Muscle-RE or in Liver+Muscle-RE, where both clocks potentially work in synergy. Even though liver and muscle transcriptional responses to night feeding were similar in single versus double RE mice, if an important clock connection does indeed exist along this axis, then double RE mice would show a substantial improvement in glucose tolerance over single RE mice. Indeed, under night feeding, oral glucose tolerance tests revealed a synergistic effect of having both liver and muscle clocks (Fig. 4f,g). Strikingly, Liver+Muscle-RE and WT blood glucose curves were indistinguishable; 30 min post gavage, the change in blood glucose reached ~75-90 mg/dL in KO and Muscle-RE, yet just ~40 mg/dL in Liver+Muscle-RE and WT mice (Fig. 4f,g). Whilst Muscle-RE performance was improved compared to KO, this effect was not statistically significant. These results demonstrate a functional connection between liver and muscle clocks for systemic glucose metabolism in the presence of a feeding-fasting rhythm (Fig. 4h).

In summary, we reveal tissue-specific differences in the autonomous capacities of peripheral clocks, showing that the muscle clock is less autonomous than the liver clock regarding both the robustness of core clock oscillations and extent of rhythmic transcriptional output. Combined with our reports on liver and skin (*4*, *5*, *7*, *8*, *25*), this reveals a differential dependence of each peripheral tissue clock on inputs from the rest of the clock network for full transcriptional function (*26*). Even though rhythmic transcriptional output of the autonomous muscle clock is low, we still find restoration of aspects of glucose metabolism, revealing a core autonomous function of the muscle clock and supporting previous findings from muscle-specific Bmal1-KO mice which demonstrated necessity of the muscle clock for efficient glucose uptake (*12*, *13*). Similar to our previous findings in Liver-RE mice (*8*), we also find evidence that muscle *Bmal1* acts as an integrator of feeding-fasting signals to support core clock and improvements in output gene expression. For mice expressing either liver or muscle autonomous clocks alone, however, the addition of feeding rhythm was not sufficient for functional restoration of glucose tolerance. Both clocks were required in combination with feeding-fasting for glucose tolerance akin to wild type. The mechanism of action for this rescue remains an open question. Our previous work has implicated soluble factors, as-of-yet unidentified, in signaling between the muscle clock and liver (*8*). Indeed, the liver and muscle are in a state of shared metabolic flux under conditions of feeding and fasting (*15*, *17*), supporting energy production, storage and usage within each tissue. From our data, we conclude that diurnal regulation of glucose pathways stemming from both liver and muscle clocks, along with the feeding-fasting rhythm, are required for the efficient uptake and processing of glucose and glucose-derived substrates from the circulation.

In this study, the use of *Bmal1* reconstitution mice allowed us to address questions of sufficiency within the clock network and identify conditions of peripheral-to-peripheral clock connections. One limitation of these mice is that overall systemic metabolism is shaped by a mutant hypothalamus and other *Bmal1*-null metabolic organs (*4*, *5*, *7*, *9*, *25*). As a result, their systemic metabolic state is different than wildtype, even under conditions of night feeding (*8*). Still, we show here and have shown previously that reconstituted peripheral clocks can overcome this altered context, especially with the addition of night feeding (*8*). For example, though pancreas BMAL1 regulates glucose-stimulated insulin secretion from beta cells and is a key peripheral node of the clock system with respect to feeding-fasting behavior (*27*, *28*), night feeding of Liver+Muscle-RE mice induced a serum insulin rhythm and rescued systemic glucose tolerance even in the absence of pancreas BMAL1. In addition, central clock reconstituted mice also exhibit improved glucose tolerance, at least in part due to reinstated feeding-fasting behavior (*7*). Considering then all evidence, it is likely that in the wildtype setting, liver, muscle and pancreas clocks, as well as the endogenous feeding rhythm set by hypothalamic clocks, constitute layers of regulation on daily systemic glucose homeostasis that work in concert. Additional work is now required to explore the impact of liver clock-muscle clock connections under physiological and pathophysiological states, and to assess the impact of additional tissue clocks on this axis.

Overall, this study highlights that circadian clock programming inherent to liver and muscle is not sufficient for a glucose tolerance, a systems level function that requires contributions from the two organs. Rather, additional inputs from other clocks, such as the feeding-fasting rhythm controlled by the central clock, are required for metabolic synergy between these two peripheral clocks. This work demonstrates a minimal circadian clock network for glucose tolerance, revealing inter-tissue clock communication as a key component supporting systemic metabolic homeostasis.

## Methods

### Animals

Mice were bred and housed at the University of California Irvine vivarium, in accordance with the guidelines of the Institutional Animal Care and Use Committee. Animal experiments were performed while considering ARRIVE (Animal Research: Reporting of In Vivo Experiments) guidelines, as follows. *Bmal1*-stop-FL mice were generated on the C57BL/6J background as previously described (*1*) (see also Fig. 1). Liver reconstituted (Liver-RE; hepatocyte-specific reconstitution of *Bmal1*) mice were generated by crossing *Bmal1*-stop-FL with Alfp-Cre mice as previously described (*2*). Skeletal muscle reconstituted (Muscle-RE; myocyte-specific reconstitution of *Bmal1*) mice were generated by crossing *Bmal1*-stop-FL with mice expressing *cre* recombinase driven by the human alpha-skeletal actin (Hsa; *Acta1*) promotor (Hsa-Cre). Crosses of *Bmal1*-stop-FL with both Cre lines generated Liver+Muscle reconstituted (Liver+Muscle-RE) mice. Experimental genotypes were: WT – *Bmal1*^wt/wt^, Alfp-Cre^-/tg^, Hsa-Cre^-/tg^; KO – *Bmal1*^stop-FL/stop-FL^, Alfp-Cre^-/-^, Hsa-Cre^-/-^; Liver-RE – *Bmal1*^stop-FL/stop-FL^, Alfp-Cre^-/tg^, Hsa-Cre^-/-^; Muscle-RE – *Bmal1*^stop-FL/stop-FL^, Alfp-Cre^-/-^, Hsa-Cre^-/tg^; Liver+Muscle-RE – *Bmal1*^stop-FL/stop-FL^, Alfp-Cre^-/tg^, Hsa-Cre^-/tg^. Alfp-Cre and Hsa-Cre positive WTs were used as controls unless otherwise indicated. All experiments used 8- to 12-week-old mice entrained to a 12 hr light: 12 hr dark cycle and fed *ad libitum* or placed under night feeding (NF) wherein mice were given *ad libitum* access to food during the dark period from zeitgeber time (ZT)12 to ZT24/0. For feasibility purposes, males were used for transcriptomic analyses and females for functional experiments. Key functional experiments were confirmed in both male and female mice.

### Western Blot

Frozen tissue was homogenized in lysis buffer (RIPA - 50 mM Tris-HCl [pH 8], 150 mM NaCl, 5 mM EDTA, 15 mM MgCl2 and 1% NP-40) supplemented with Protease Inhibitor Cocktail (Roche, Basel, Switzerland), 500 μM PMSF (serine protease inhibitor), 10 mM nicotinamide (Sirtuin inhibitor) and 330 nM TSA (Class I and II HDAC inhibitor). Samples were lysed on ice with periodic mixing for 30 min. Next, samples were sonicated (5 sec on, 5 sec off, 4 cycles, 60% amp.) and centrifuged at 13,200 rpm at 4°C for 15 min and the supernatant was collected. We used the Braford method to determine protein concentration, using Protein Assay Dye (BioRad). 20 μg protein from each sample was separated on 6% or 8% gels by SDS-PAGE and subsequently transferred to a nitrocellulose membrane. Blots were then blocked with 5% milk in TBS-T (0.1% Tween-20, TBS) at room temperature for 2 hr. Primary antibodies were diluted in 5% milk TBS-T and incubated with blots at 4°C overnight (BMAL1, Abcam – ab93806; ACTIN, Abcam – ab3280, P84, Genetex – GTX70220). Blots were incubated with HRP-conjugated secondary antibodies (Mouse IgG-HRP conjugate, EMD Millipore – AP160P; Rabbit IgG-HRP linked, EMD Millipore – 12–348) at room temperature for 1 hr and visualized with Immobilon Western chemiluminescent HRP substrate (Millipore, Burlington, MA). Finally, blots were developed on HyBlot CL autoradiography film (Denville Scientific, Holliston, MA).

### Indirect Calorimetry and Food Intake

Metabolic parameters were measured using the Phenomaster metabolic cage system (TSE Systems Inc., Bad Homburg, Germany). The climate chamber was set to 21°C and 50% humidity with a 12h:12h light–dark cycle. Mice were maintained on standard chow (Teklad 2020x) *ad libitum* or time-restricted as indicated, singly housed, and acclimated for 48h prior to data collection. VO2, VCO2, respiratory exchange ratio (RER), energy expenditure and food intake were monitored for 3 min per cage and measured every 39 min for four to six consecutive days. Body composition (lean mass and fat mass) was measured in non-anesthetized mice via rodent MRI (QNMR EchoMRI100; Echo Medical Systems) prior to recordings for normalization of data. Food intake values were calculated by multiplying average grams eaten per hour by the kcal/g of the chow diet.

### Locomotor Activity

Mice were individually housed in home cages equipped with optical beam motion detection capability (Starr Life Sciences). Data were collected using VitalView 5.0 software and analyzed using Clocklab software (Actimetrics).

### RNA-Sequencing

Total RNA was monitored for quality control using the Agilent Bioanalyzer Nano RNA chip and Nanodrop absorbance ratios for 260/280nm and 260/230nm. Library construction was performed according to the Illumina TruSeq Total RNA stranded protocol. The input quantity for total RNA was 850ug and rRNA was depleted using ribo-zero rRNA gold removal kit (human/mouse/rat). The rRNA-depleted RNA was chemically fragmented for three minutes. First strand synthesis used random primers and reverse transcriptase to make cDNA. After second strand synthesis the cDNA was purified using AMPure XP beads, the cDNA was end-repaired and the 3’ ends were adenylated. Illumina unique dual indexed adapters were ligated on the ends and the adapter-ligated fragments were enriched by nine cycles of PCR. The resulting libraries were validated by qPCR and sized by Agilent Bioanalyzer DNA high sensitivity chip. The concentrations for the libraries were normalized and then multiplexed together. The multiplexed libraries were sequenced using paired-end 100 cycles chemistry on the NovaSeq 6000.

### RNA-Sequencing Analysis

Sequencing results arrived in FastQ format. The pair-end reads from each replicate experiment were aligned to the reference genome assembly mm10 and the corresponding transcriptome using the Tuxedo protocol [1]. Reads uniquely aligned to known exons, or splice junctions, extracted with no more than two mismatches were included in the transcriptome. Gene expression levels were directly computed from the read alignment results for each replicate. Standard FPKM (fragments per kilobase of exon per million mapped reads) values were extracted for each gene covered by the sequencing data and each replicate used in this study. Venn diagrams were created using Biovenn (*3*) followed by processing with Adobe Illustrator for display.

### Rhythmicity and Gene Set Enrichment Analysis

To identify oscillating transcripts, transcriptomic data for each tissue was processed via JTK_CYCLE (*4*) (ADJ P<0.01, 24-hour period) using R-studio. We used BIO_CYCLE (p<0.01) as an additional measure of rhythmicity (*5*). Gene set enrichment analysis using KEGG pathways was performed with DAVID.

### Oral Glucose Tolerance Test

Mice were fasted from ZT0 to ZT6 and gavaged with 1.5 g/kg glucose. Blood glucose was measured via the tail at 0, 15, 30, 60 and 120 min following the gavage using an Accu-Chek Aviva Plus Blood Glucose Monitoring System. Glucose tolerance curves were calculated using current standards (*6*); baseline was subtracted from raw blood glucose values at each time point to give delta or change in glucose concentration over time. Area under the curve was calculated using standard parameters.

### Glucose Tracing

Glucose tracing was performed identically to oral glucose tolerance tests, except gavaged glucose was uniformly labeled with carbon 13 (U-^13^C_6_, Cambridge Isotope Laboratories, CLM-1396-PK). Serum and tissues were harvested 25-30 min after gavage.

### qPCR

For liver and muscle, 1μg of Trizol-extracted RNA was reverse transcribed using Thermo Scientific Maxima Reverse Transcriptase and qPCR performed using Syber Green Master Mix on a ThermoFisher Quantstudio 3. All gene expression data normalized to 18S ribosomal RNA. Primer sequenced used:

BMAL1 F GCAGTGCCACTGACTACCAAGA

BMAL1R TCCTGGACATTGCATTGCAT

Rn18SF CGCCGCTAGAGGTGAAATTC

Rn18SR CGAACCTCCGACTTTCGTTCT

### Tracing sample preparation and metabolite extraction

Serum (5 μl) was mixed with 150 μl 4°C 40:40:20 methanol:acetonitrile:water (extraction solvent), vortexed and immediately centrifuged at 16,000g for 10 min at 4°C. The supernatant (~ 100 μl) was collected for liquid chromatography-mass spectrometry (LC-MS) analysis. Frozen tissue samples were ground at liquid nitrogen temperature with a CryoMill (Retsch). To minimize data variation due to tissue heterogeneity, entire tissues were collected and ground. The resulting tissue powder (approximately 20 mg) was weighed and then extracted by adding 4°C extraction solvent, vortexed and centrifuged at 16,000g for 10 min at 4°C. The volume of the extraction solution (μl) was 40x the weight of tissue (mg) to make an extract of 25 mg of tissue per ml of solvent. The supernatant (40 μl) was collected for LC-MS analysis.

### Metabolite measurements using LC-MS

A quadrupole orbitrap mass spectrometer (Q Exactive; Thermo Fisher Scientific) operating in negative or positive ion mode was coupled to a Vanquish UHPLC system (Thermo Fisher Scientific) with electrospray ionization and used to scan from m/z 70 to 1,000 at 1 Hz, with a 140,000 resolution. LC separation was achieved on an XBridge BEH Amide column (2.1 x 150 mm^2^, 2.5 μm particle size, 130 Å pore size; Waters Corporation) using a gradient of solvent A (95:5 water: acetonitrile with 20 mM of ammonium acetate and 20 mM of ammonium hydroxide, pH 9.45) and solvent B (acetonitrile). Flow rate was 150 μl/min. The LC gradient was: 0 min, 85% B; 2 min, 85% B; 3 min, 80% B; 5 min, 80% B; 6 min, 75% B; 7 min, 75% B; 8 min, 70% B; 9 min, 70% B; 10 min, 50% B; 12 min, 50% B; 13 min, 25% B; 16 min, 25% B; 18 min, 0% B; 23 min, 0% B; 24 min, 85% B; and 30 min, 85% B. The autosampler temperature was 5°C and the injection volume was 3 μl. Data were analyzed using the MAVEN software (build 682, http://maven.princeton.edu/index.php). Natural isotope correction was performed with AccuCor R code (https://github.com/lparsons/accucor).

### Insulin Measurements

Collected blood was placed on ice and subsequently centrifuged at 3,000 rpm for 10 min at 4° C. The supernatant was stored at −80° C. All samples were thawed and used as starting material for the Ultra-Sensitive Mouse Insulin ELISA Kit (Cystal Chem 90080), which was utilized according to the manufacturer’s instructions.

### Statistics

All data are displayed as mean ± S.E.M. unless otherwise noted. For each experiment, the number of biological replicates, statistical test and significance threshold can be found in the figure legend or main text. Complex statistical analyses are described within the corresponding methods section. Data were analyzed in Prism 6.0 software (GraphPad). Sample size was determined by referencing literature standards for studies of circadian rhythms (ref).

## Acknowledgements

This paper is dedicated to the memory of Professor Paolo Sassone-Corsi, a hugely inspiring scientist and mentor and who remains an important influence on our work. We extend special thanks to our animal technician S. Sato and laboratory manager W. Orquiz for valuable help, as well as to all members of the P.S.C. laboratory for support. J.S. was supported by Zymo-CEM Postdoctoral Fellowship (Zymo Research) awarded at the University of California, Irvine. K.B.K. is supported by NIH, NIDDK F32 Fellowship - DK121425. T.S. was supported by a Japan Society for the Promotion of Science (JSPS) fellowship. C.M.G. was supported by the National Cancer Institute of the National Institutes of Health (NIH) under award number T32CA009054 and by the European Union’s Horizon 2020 research and innovation programme under the Marie Sklodowska-Curie grant agreement 749869. P.P. was funded by The Wenner-Gren Foundations, The Foundation Blanceflor Boncompagni Ludovisi, née Bildt and Tore Nilsson Foundation for Medical Science. A.V. was supported by the Hitachi-Nomura postdoctoral fellowship awarded through the Department of Biological Chemistry at the University of California, Irvine. The work of S.C., M.S., and P.B. was in part supported by NIH grant GM123558 C.J. was supported by the AASLD Foundation Pinnacle Research Award in Liver Disease, the Edward Mallinckrodt, Jr. Foundation Award, and NIH/NIAAA R01 AA029124. P.S.W. is supported by grant RYC2019-026661-I funded by MCIN/AEI/10.13039/501100011033 and by “ESF Investing in your future”. Financial support for the S.M. Laboratory is provided through the NIH/NCI (Grants: R01CA244519, R01CA259370, K22CA212045). Research in the S.A.B. lab is supported partially by the European Research Council (ERC) under the European Union’s Horizon 2020 research and innovation programme (Grant agreement No. 787041), the Government of Cataluña (SGR grant), the Government of Spain (MINECO), the La Marató/TV3 Foundation, the Foundation Lilliane Bettencourt, the Spanish Association for Cancer Research (AECC) and The Worldwide Cancer Research Foundation (WCRF). The IRB Barcelona is a Severo Ochoa Center of Excellence (MINECO award SEV-2015-0505). P.M.C. acknowledges funding from MICINN-RTI2018-096068, ERC-2016-AdG-741966, LaCaixa-HEALTH-HR17-00040, MDA, UPGRADE-H2020-825825, AFM, DPP-Spain, Fundació La MaratóTV3-80/19-202021, MWRF, and María-de-Maeztu Program for Units of Excellence to UPF (MDM-2014-0370) and the Severo-Ochoa Program for Centers of Excellence to CNIC (SEV-2015-0505). Work in the P.S.-C. laboratory supported by NIH grants R21DK114652 and R21AG053592, a Challenge Grant from the Novo Nordisk Foundation (NNF-202585), KAUST funding (OSR-2019-CRG8-URF/1/4042), and via access to the Genomics High Throughput Facility Shared Resource of the Cancer Center Support Grant (CA-62203) and the UCI and NIH-shared instrumentation grants 1S10RR025496-01, 1S10OD010794-01, and 1S10OD021718-01.

---

**Extended Data Figure 1.**
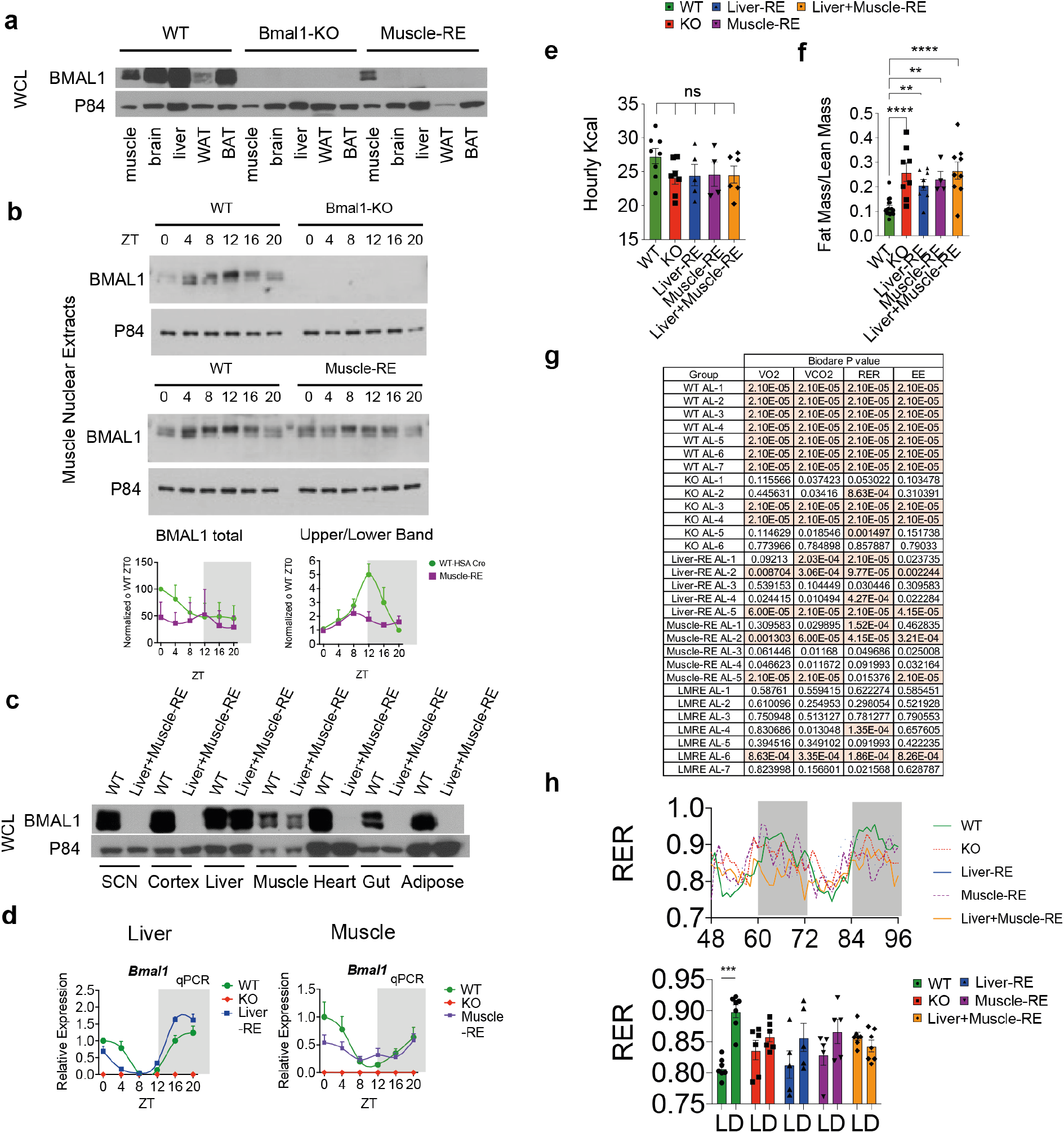
Related to Figure 1. **a**, whole-cell lysates (WCL) from various tissues collected at ZT4. WAT – white adipose tissue; BAT – brown adipose tissue. **b**, top, nuclear protein extracts from muscles collected at the indicated time points. bottom, densitometry quantification of n=3 biological replicates, normalized to p84 for BMAL1 total. **c**, WCLs from various tissues collected at ZT4 **d**, qPCR of *bmal1* expression in liver and muscle showing confirmation of knockout, n=3-6. **e**, daily food intake expressed as average kcals per hr. One-way ANOVA with Fisher’s LSD, n=4-8, ns=not significant. **f**, fat mass to lean mass ratio measured by ecoMRI. One-way (*8*)ANOVA with Fisher’s LSD, n=4-16, ** p<0.01, **** p<0.0001. **g**, rhythmicity analysis of metabolic cage parameters using BioDare2. Value shown is p-value for each parameter for individual mice, orange-highlighted cells are significant at p<0,01. VO2 – oxygen consumption rate, VCO2 – carbon dioxide release rate, EE – energy expenditure; RER – respiratory exchange ratio. **h**, Top – mean RER reading over a 48h period, of mice under 12 hr light: 12 hr dark conditions. Bottom – bar graph showing light (L) and dark (D) phase averages. Two-way ANOVA with Bonferroni post-hoc, n=5-7, *** p<0.001.

**Extended Data Figure 2.**
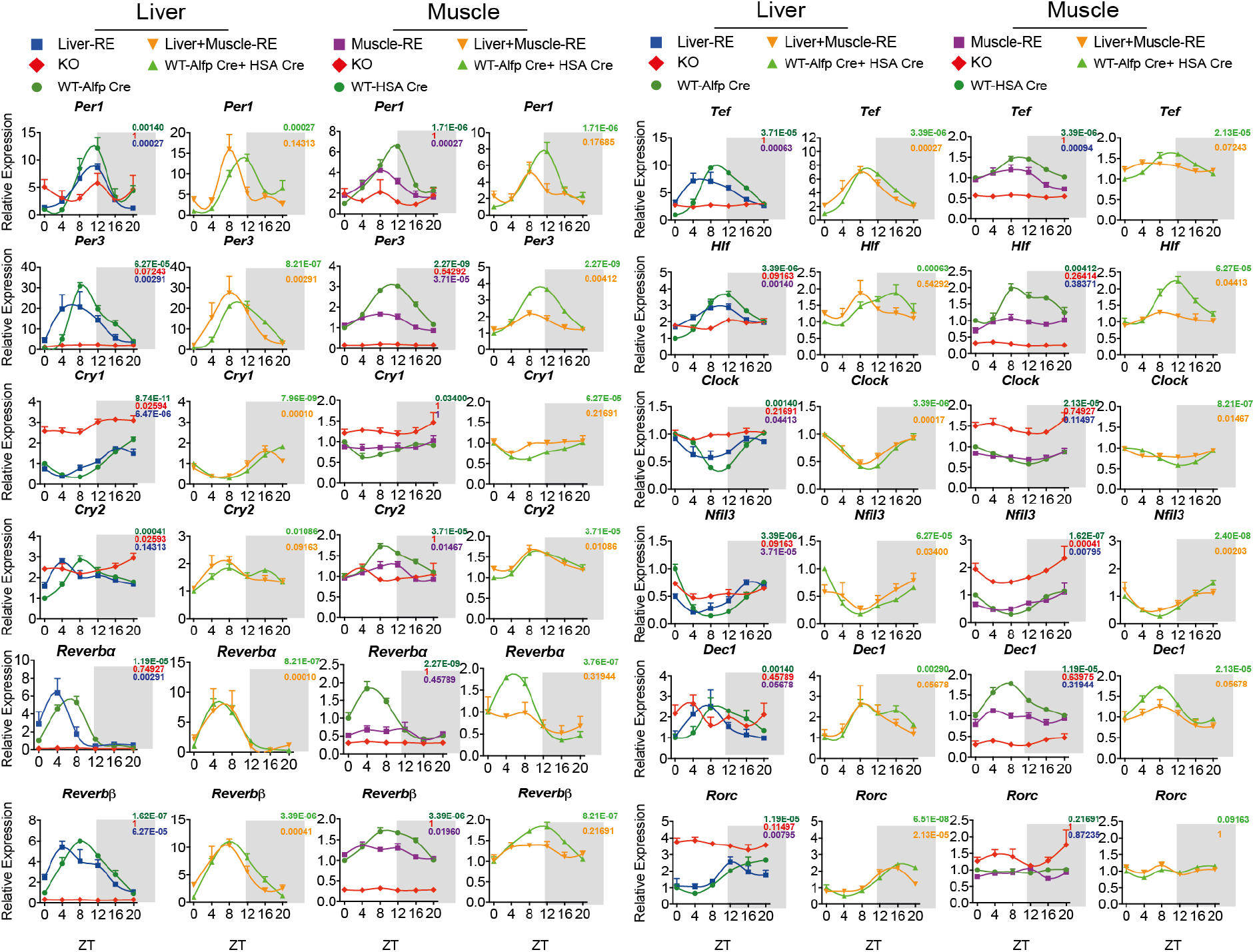
Related to Figure 1. Expression of clock genes from RNA sequencing of livers and gastrocnemius muscles harvested around the clock under 12 hr light: 12 hr dark conditions (n=3). Liver sequencing data for WT-Alfp Cre, KO and Liver-RE from (*8*). Values shown are JTK_CYCLE adjusted p-values.

**Extended Data Figure 3.**
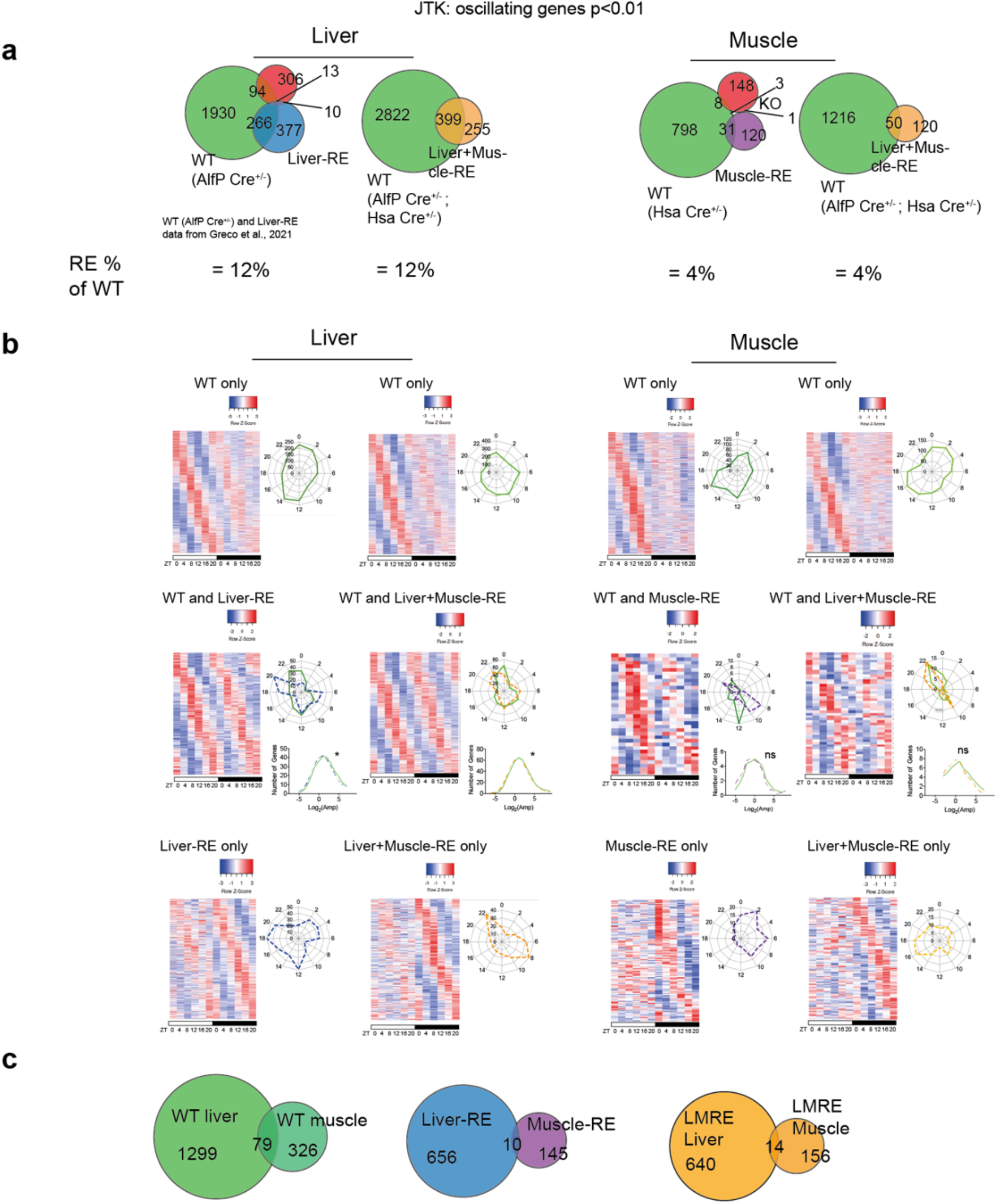
Related to Figure 2. JTK_CYCLE rhythmicity analysis of RNA sequencing of livers and gastrocnemius muscles harvested around the clock under 12 hr light: 12 hr dark conditions (n=3). Liver sequencing data for WT-Alfp Cre, KO and Liver-RE from (*8*). **a**, Venn diagrams showing overlap of oscillating genes between genotypes. **b**, left, phase sorted heatmaps of genes oscillating in WT and RE mice; right-top, phase of oscillating genes; right-bottom, amplitude of oscillating genes. Mann-Whitney test, ns=not significant, * p<0.05, n=3. **c**, overlap of oscillating genes between liver and muscle. For WT groups, genes oscillating in both cre backgrounds were included.

**Extended Data Figure 4.**
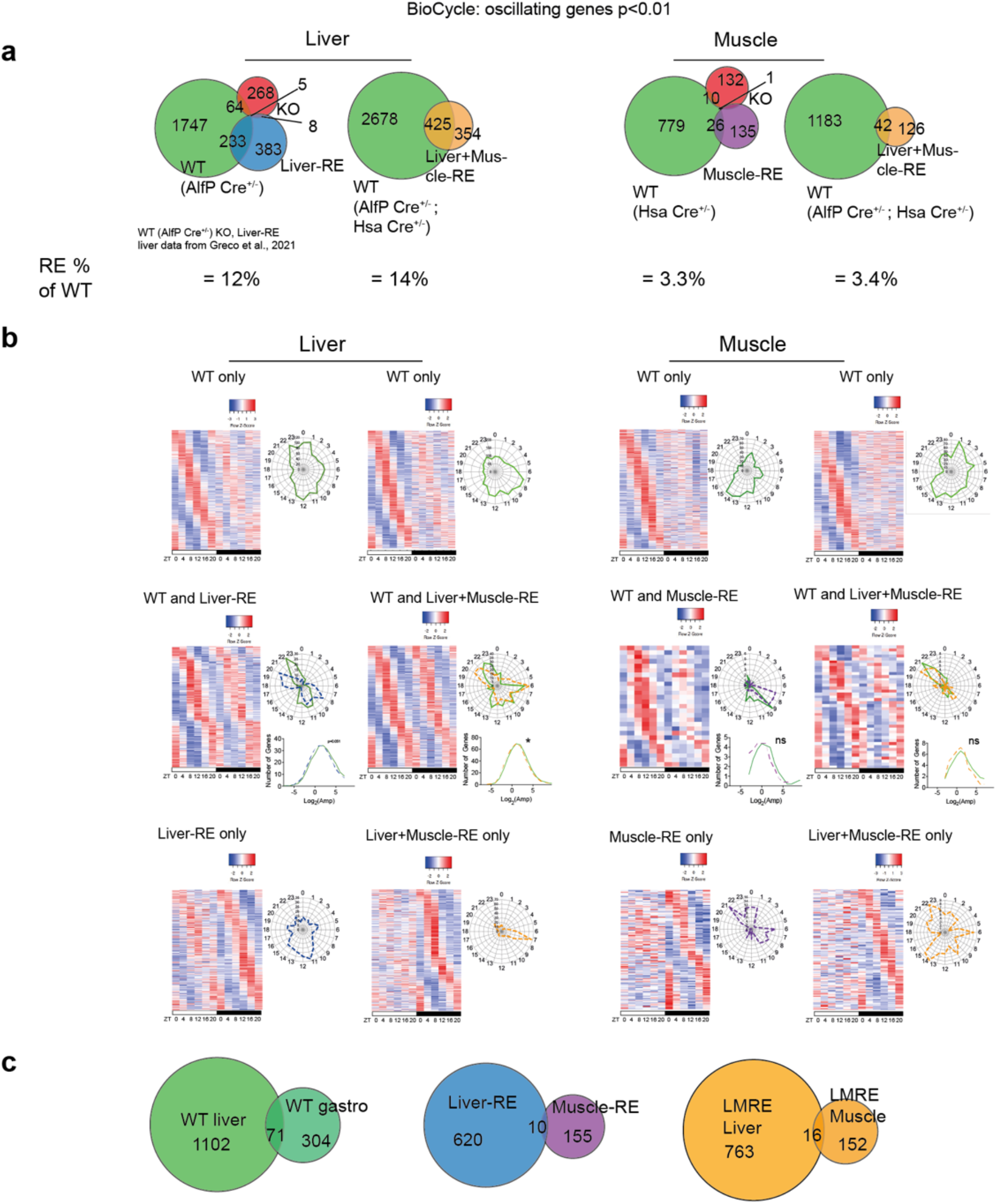
Related to Figure 2. BIO_CYCLE rhythmicity analysis of RNA sequencing of livers and gastrocnemius muscles harvested around the clock under 12 hr light: 12 hr dark conditions (n=3). Liver sequencing data for WT-Alfp Cre, KO and Liver-RE from (*8*). **a**, Venn diagrams showing overlap of oscillating genes between genotypes. **b**, left, phase sorted heatmaps of genes oscillating in WT and RE mice; right-top, phase of oscillating genes; right-bottom, amplitude of oscillating genes. Mann-Whitney test, ns=not significant, * p<0.05, n=3. **c**, overlap of oscillating genes between liver and muscle. For WT groups, genes oscillating in both cre backgrounds were included.

**Extended Data Figure 5.**
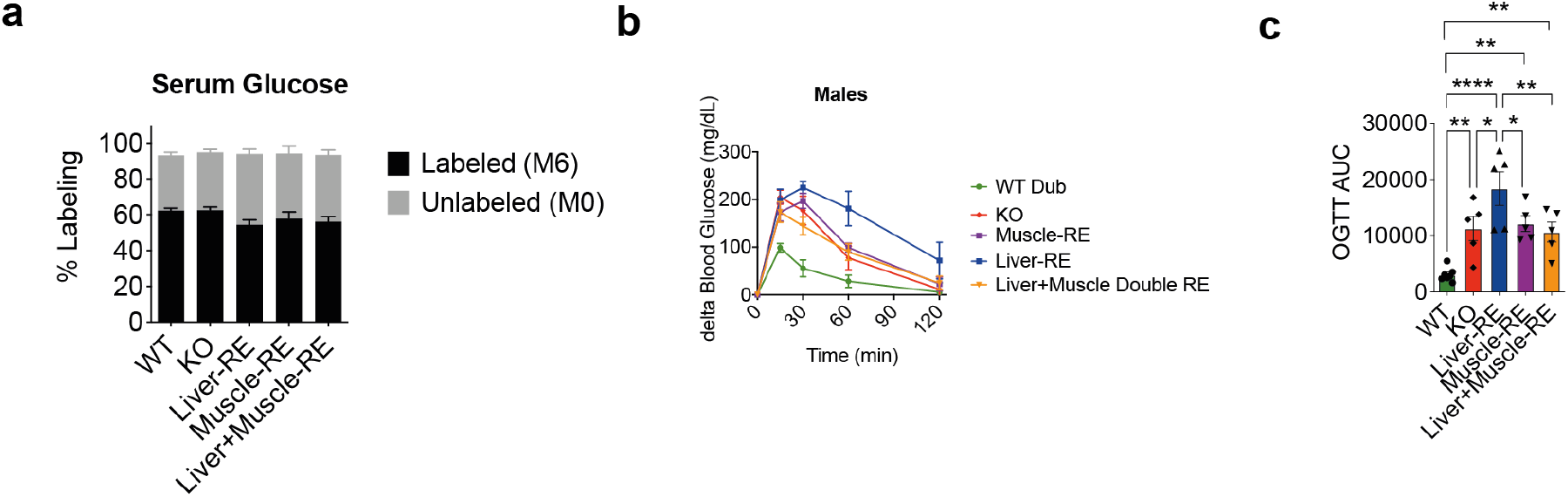
Related to Figure 3. **a**, Percentage of uniformly labeled (M6) vs unlabeled (M0) glucose in serum 25-30 min following oral gavage showing consistent enrichment of labeled glucose in the blood across genotypes. **b**, oral glucose tolerance tests (OGTT) were performed on male mice as described in Fig. 3 (n=6-7) and are expressed as delta change from baseline blood glucose. **c**, area under the curve (AUC) for OGTTs, * p<0.05, ** p<0.01, *** p<0.001, one-way ANOVA with Fishers LSD test.

**Extended Data Figure 6.**
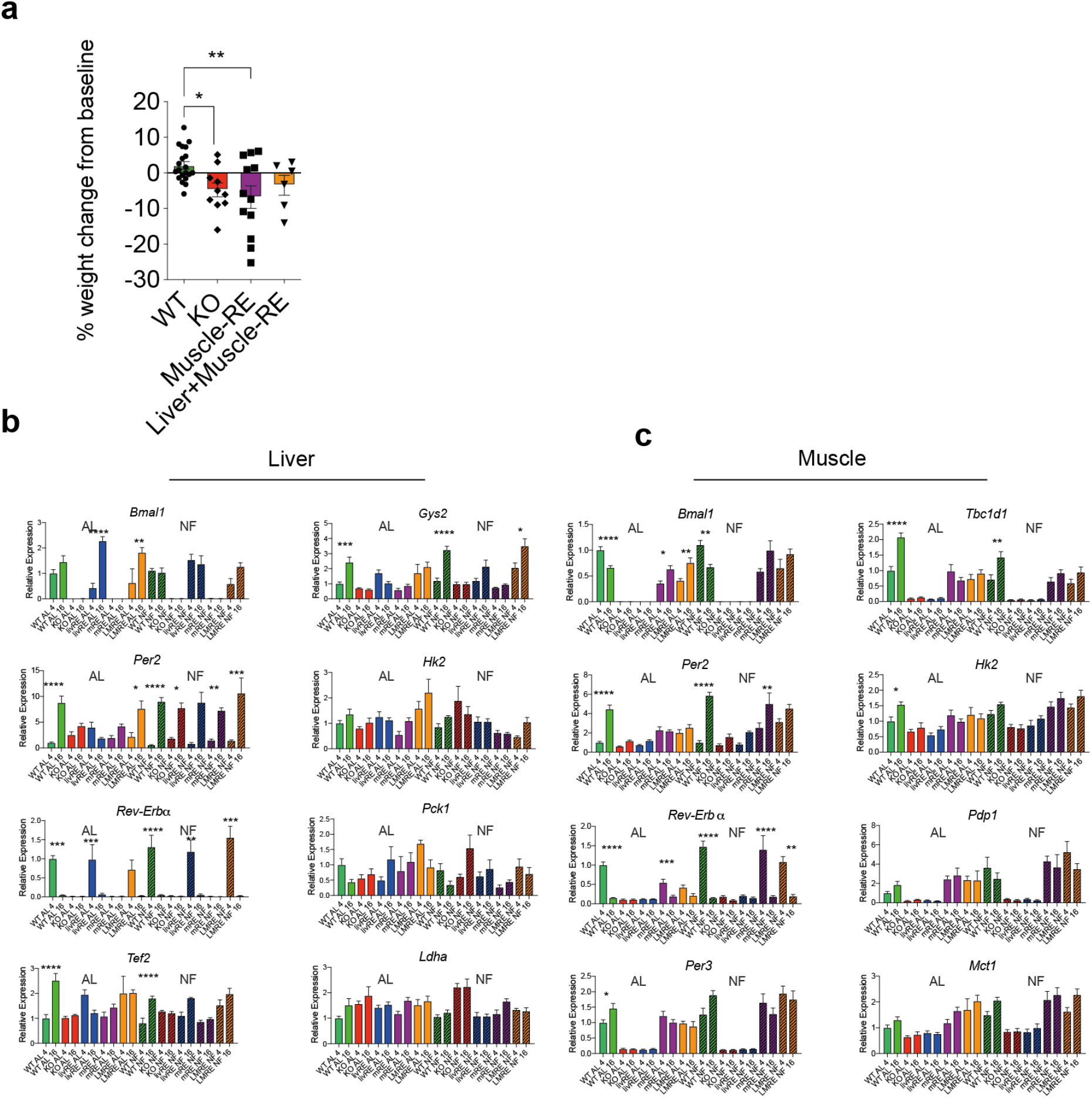
Related to Figure 4. **a**, weight change from baseline taken during *ad libitum* feeding to after 2 weeks of night feeding. One-way ANOVA with Fisher’s LSD, * p<0.05, ** p<0.01, n=6-20. **b-c**, gene expression by qPCR in liver and muscle of clock and metabolic genes at ZT4 and 16 under AL and NF conditions, data same as in Figure 4-except with the addition of single RE controls in opposing tissues; liver-RE for muscle data, and muscle-RE for liver data n=3-6. * p<0.05, ** p<0.01, *** p<0.001, ****p<0.0001 by two-way ANOVA with Bonferroni post-test.

